# Edge-centric analysis of time-varying functional brain networks with applications in autism spectrum disorder

**DOI:** 10.1101/2021.07.01.450812

**Authors:** Farnaz Zamani Esfahlani, Lisa Byrge, Jacob Tanner, Olaf Sporns, Daniel P. Kennedy, Richard F. Betzel

## Abstract

The interaction between brain regions changes over time, which can be characterized using time-varying functional connectivity (tvFC). The common approach to estimate tvFC uses sliding windows and offers limited temporal resolution. An alternative method is to use the recently proposed edge-centric approach, which enables the tracking of moment-to-moment changes in co-fluctuation patterns between pairs of brain regions. Here, we first examined the dynamic features of edge time series and compared them to those in the sliding window tvFC (sw-tvFC). Then, we used edge time series to compare subjects with autism spectrum disorder (ASD) and healthy controls (CN). Our results indicate that relative to sw-tvFC, edge time series captured rapid and bursty network-level fluctuations that synchronize across subjects during movie-watching. The results from the second part of the study suggested that the magnitude of peak amplitude in the collective co-fluctuations of brain regions (estimated as root sum square (RSS) of edge time series) is similar in CN and ASD. However, the trough-to-trough duration in RSS signal is greater in ASD, compared to CN. Furthermore, an edge-wise comparison of high-amplitude co-fluctuations showed that the within-network edges exhibited greater magnitude fluctuations in CN. Our findings suggest that high-amplitude co-fluctuations captured by edge time series provide details about the disruption of functional brain dynamics that could potentially be used in developing new biomarkers of mental disorders.

## INTRODUCTION

The human brain is fundamentally a complex system and can be modeled as a network of functionally connected brain regions [1, 2]. In practice, functional connectivity (FC) is estimated as the Pearson correlation of brain regions’ functional magnetic resonance imaging (fMRI) blood oxygen level-dependent (BOLD) time courses, often recorded in the absence of explicit task instructions, i.e. the resting state [3, 4]. A growing number of studies have used FC to link inter-individual variation in brain network organization with cognition [5], development [6], and disease [7].

In most applications, FC is estimated using data from an entire scan session, resulting in a single connectivity matrix whose weights express the average connection strength between pairs of brain regions [8]. However, the brain’s macro-scale functional organization varies over shorter timescales on the order of seconds [9–11]. To capture these changes, many studies have estimated FC over shorter intervals using dynamic or time-varying FC (tvFC) [12]. In most cases, tvFC is estimated using a sliding window method. In this approach, FC is estimated using only frames that fall within a window of fixed duration. The window is advanced by some amount, and the process repeated. In the end, the result is a sequence of FC estimates.

Sliding window time-varying FC (sw-tvFC) has been used widely in order to characterize time-varying changes in brain network organization in general, but also to study how fluctuations in brain network architecture accompany cognitive processes across time [13, 14]. In addition, tvFC has proven useful for generating novel biomarkers [13, 15–17].

Despite its success and continued application, sliding-window methods have a number of limitations. First, they require users to choose a series of parameters, including window duration, shape, and the amount of overlap between successive windows [18–21]. These decisions are non-trivial and, in general, impact the inferred patterns of connectivity. They can also introduce artifacts into estimates of time-varying FC, e.g., through aliasing effects. Perhaps most serious, sliding window methods make it impossible to precisely localize changes in FC to a specific instant in time. The very nature of a window means that FC receives contributions from all points within that interval. Collectively, these limitations present challenges, both in estimating and interpreting time-varying FC estimated using sliding window techniques [22, 23].

Recently, we proposed a novel, edge-centric method for estimating time-varying FC [24, 25]. This method precisely decomposes FC into its framewise contributions, yielding a frame-by-frame account of interregional co-fluctuations across time, which we refer to as cofluctuation or edge time series (ETS). A key feature of this approach is that ETS are estimated without specifying parameters or the need to perform any windowing. Consequently, many of the limitations associated with sliding window methods are not applicable. Since its introduction, ETS has been used to study individual differences [26] and the origins of brain systems [27], and its anatomical underpinnings examined using *in silico* models [28]. However, the performance of ETS has not been systematically compared with that of sliding window techniques. Additionally, because edge time series represent a new construct, their utility for linking brains with behavior remains unclear.

Here, we address these gaps in knowledge. In the first section of the paper, we conduct a systematic comparison of temporal properties of ETS and sw-tvFC. Our findings show two main features of ETS that can not be seen in sw-tvFC. First, ETS exhibits rapid and bursty fluctuations at rest, as evidenced by reduced autocorrelation and more frequent transitions from one brain state to another. In addition, these co-fluctuations were synchronized across subjects during movie-watching condition. Second, collective fluctuations of ETS showed less similarity between their high and low-amplitude, which was indicated by higher peak amplitudes and shorter trough-to-trough duration (number of frames between two local minima) compared to sw-tvFC. Building on these two important features of ETS, in the second part of the paper, we studied differences in the co-fluctuations of brain regions between autism spectrum disorder (ASD) and healthy control (CN) subjects during movie-watching condition. Our findings suggested that overall, the peak amplitude of collective co-fluctuations of brain regions is similar in ASD and CN, however the trough-to-trough duration is greater in ASD. Additionally, a detailed analysis of individual ETS suggested that compared to ASD, within-network edges showed higher peak co-fluctuations in CN.

## RESULTS

We applied ETS and sw-tvFC to fMRI data of 29 CN and 23 ASD subjects that were collected multiple times in resting state and during passive movie-watching conditions [29]. The overall procedures for estimating ETS and sw-tvFC and their differences are shown in Figure 1. After estimating ETS and sw-tvFC, first, in *Comparison of edge time series and sliding window-tvFC*, we used data from the CN group and compared the properties of ETS with sw-tvFC, including whole-brain co-fluctuation dynamics, synchronization of these co-fluctuations across subjects, and relationship between high and low-amplitude edge fluctuations. Next, in *Edge time series in autism spectrum disorder*, we used ETS to examine differences in the co-fluctuation patterns of brain regions in ASD and CN groups.

**FIG. 1.**
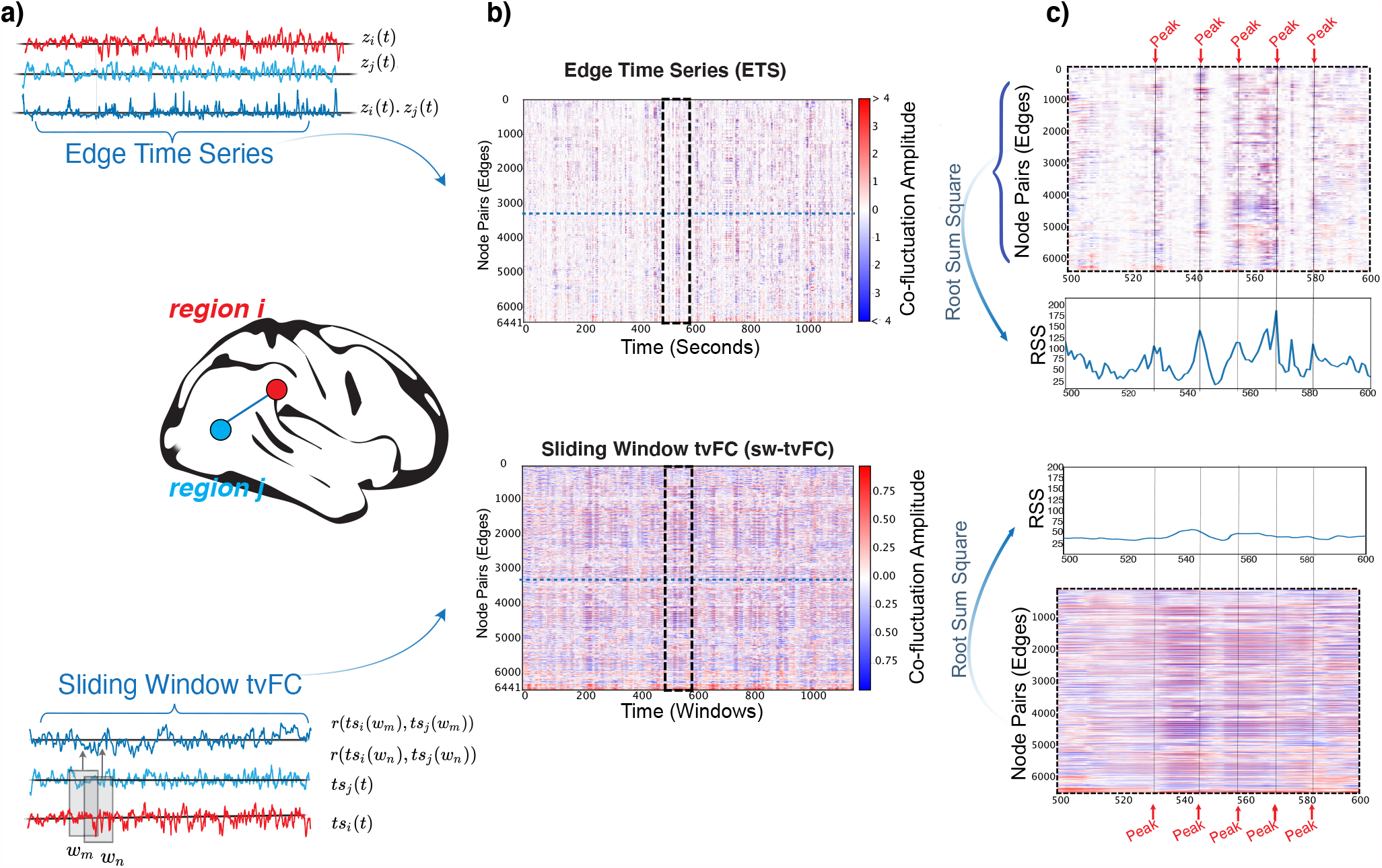
The general procedure for constructing sliding window time varying FC (sw-tvFC) networks and edge time series (ETS). (*a*) ETS are calculated as the dot product of activity of two nodes, while in the sw-tvFC, first time series is divided into equal parts (windows) and edges are estimated by calculating the correlation between time samples within each window. (*b*) After calculating ETS for all pairs of brain regions, we obtain 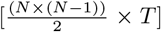 matrix. For ETS, this matrix provides detailed picture of the moment-to-moment co-fluctuations between all pairs of brain regions, whereas for sw-tvFC, this estimation is blurry due to the windowing procedure. (*c*) Whole-brain co-fluctuations can be estimated as root sum square (RSS) of all the edges fluctuations at every given time point. In ETS, the high-amplitude co-fluctuations are captured more precisely compared to sw-tvFC.

### Comparison of edge time series and sliding window-tvFC

#### Whole-brain co-fluctuation dynamics

To examine differences in the global properties of ETS and sw-tvFC, we first asked how similar are the whole-brain co-fluctuation patterns estimated by these two methods? To answer this question, we calculated ETS and sw-tvFC for every subject based on their resting-state fMRI BOLD time series. Then we vectorized the complete set of time-varying edge weights and, after resampling to ensure that ETS and sw-tvFC estimates contained the same numbers of time points, we vectorized the entire edge by time matrix and computed the similarity between conditions (Figure 2a). We repeated this procedure for sw-tvFC constructed using window sizes ranging from 10 to 100 frames in increments of 10 (each frame = 0.813 seconds). We found that sw-tvFC and ETS were moderately correlated (*r* = 0.35; window size = 20; details for other window sizes can be found in Figure 2a) suggesting that while these two methods broadly capture similar patterns of co-fluctuations, there remains considerable amounts of unexplained variance. The results for individual scans are available in Figure S1. We also explored why the correspondence between ETS with sw-tvFC was peaked at an intermediate window size (see Figure S2).

**FIG. 2.**
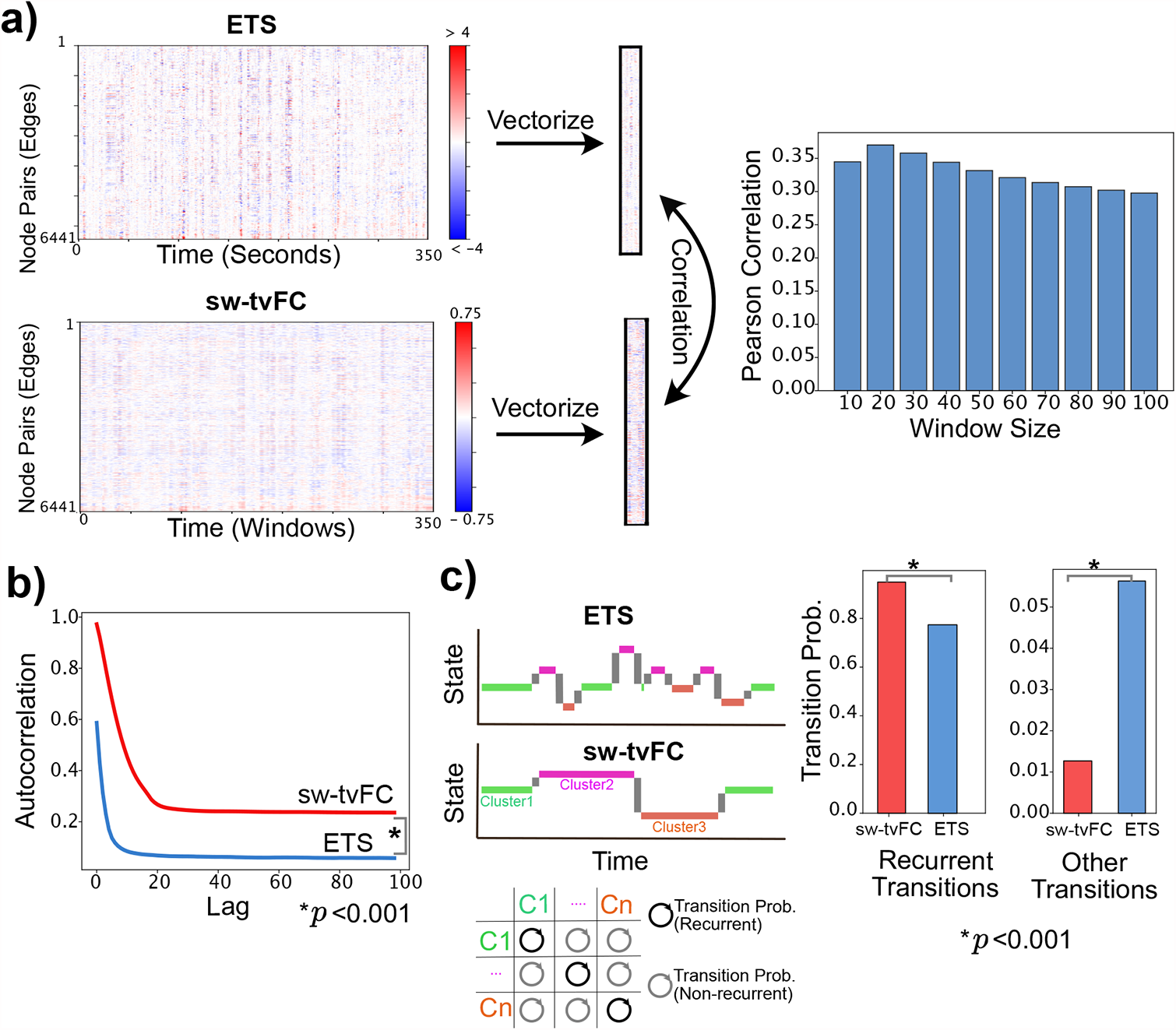
Relationship between edge time series (ETS) and sliding window-time varying functional connectivity (sw-tvFC). *(a)* ETS and sw-tvFC are calculated for every subject in resting state. ETS and sw-tvFC are resampled to ensure they contain the same numbers of time points. Then, the complete set of time-varying edge weights estimated by these two methods are vectorized and their similarity are computed. Bar plot represents the averaged Pearson correlation between all the edges in ETS and sw-tvFC across subjects. ETS and sw-tvFC are most similar for window sizes of 20 and 30. *(b)* Shows the autocorrelation and transition probabilities in ETS and sw-tvFC (i.e., window size (*w*) = 20). The autocorrelation of whole-brain co-fluctuations over time shows the lower rate of autocorrelation in ETS, suggesting the presence of high-amplitude co-fluctuations. *(c)* ETS show higher between-state transitions and lower within-state transitions compared to the sw-tvFC. States at each given time point were defined based on the whole-brain co-fluctuations.

Next, we asked to what extent time-varying changes estimated by ETS and sw-tvFC are smooth and slow *versus* abrupt and fast? To answer this question, we calculated the autocorrelation in ETS/sw-tvFC as the similarity of whole-brain co-fluctuation patterns at time *t* with the patterns at times *t* + 1, *t* + 2, …, *t* + 99, *t* + 100. Our results showed that ETS exhibited reduced averaged autocorrelation across subjects and scans compared to sw-tvFC, suggesting the presence of rapid and bursty network-level fluctuations (*t*-test, *p* < 0.001; Figure 2b; results for individual scans are in Figure S3).

Along the same line, we also asked to what extent there is a memory of previous network states in sw-tvFc versus ETS? To answer this question, we used the *k*-means clustering algorithm to cluster the time frames into non-overlapping clusters based on the similarity of whole-brain co-fluctuation patterns at different points in time [15, 30] (here we report results for *k*=5 and using subjects from all the scans; results are qualitatively similar for other values of *k* shown in Figure S3). We used these clusters to estimate the transition probabilities between all pairs of brain states, finding that ETS transitioned from one brain state to another more frequently than sw-tvFC (*t*-test, *p* < 0.001; Figure 2b; results for individual scans are in Figure S3).

Collectively, these results suggest that compared to sw-tvFC, ETS capture distinct patterns of co-fluctuations between brain regions. Our results also suggest that ETS capture a faster and more “bursty” network dynamics, in which network states change abruptly, more frequently, and over faster timescales compared to sw-tvFC. Further, these results are in line with the hypothesis that the use of sliding windows may induce smoothness in network trajectories across time, possibly obscuring rapid reconfigurations of the network over short timescales.

#### Synchronization of the whole-brain co-fluctuation patterns across subjects

In the previous section, we examined the presence of rapid and bursty fluctuations in ETS, highlighting this property as one of its main ways it differs from sw-tvFC. These high-amplitude fluctuations – referred to as “events” in previous papers [25] – are infrequent and, in previous work, were shown to be uncorrelated with in-scanner head motion [25, 26]. Therefore, they may be important in providing insights into the ongoing cognitive processes at rest and movie-watching conditions.

In this section, we examined how well co-fluctuation patterns captured by these two methods are synchronized across subjects. To address this question, we calculated the inter-subject similarity based on the collective co-fluctuations of brain regions during rest and movie-watching conditions. More specifically, the collective co-fluctuations of brain regions were estimated as the root sum square (RSS) of co-fluctuations between all pairs of brain regions (edges) at every given time point. We found that compared to sw-tvFC, the collective co-fluctuation patterns in ETS (specifically those with high-amplitude) were aligned across subjects during movie-watching condition verus resting state, (Figure 3.a). This was indicated by higher inter-subject similarity in ETS, compared to sw-tvFC during movie-watching condition (*t*-test; *p* < 0.001; Figure 3b-c).

**FIG. 3.**
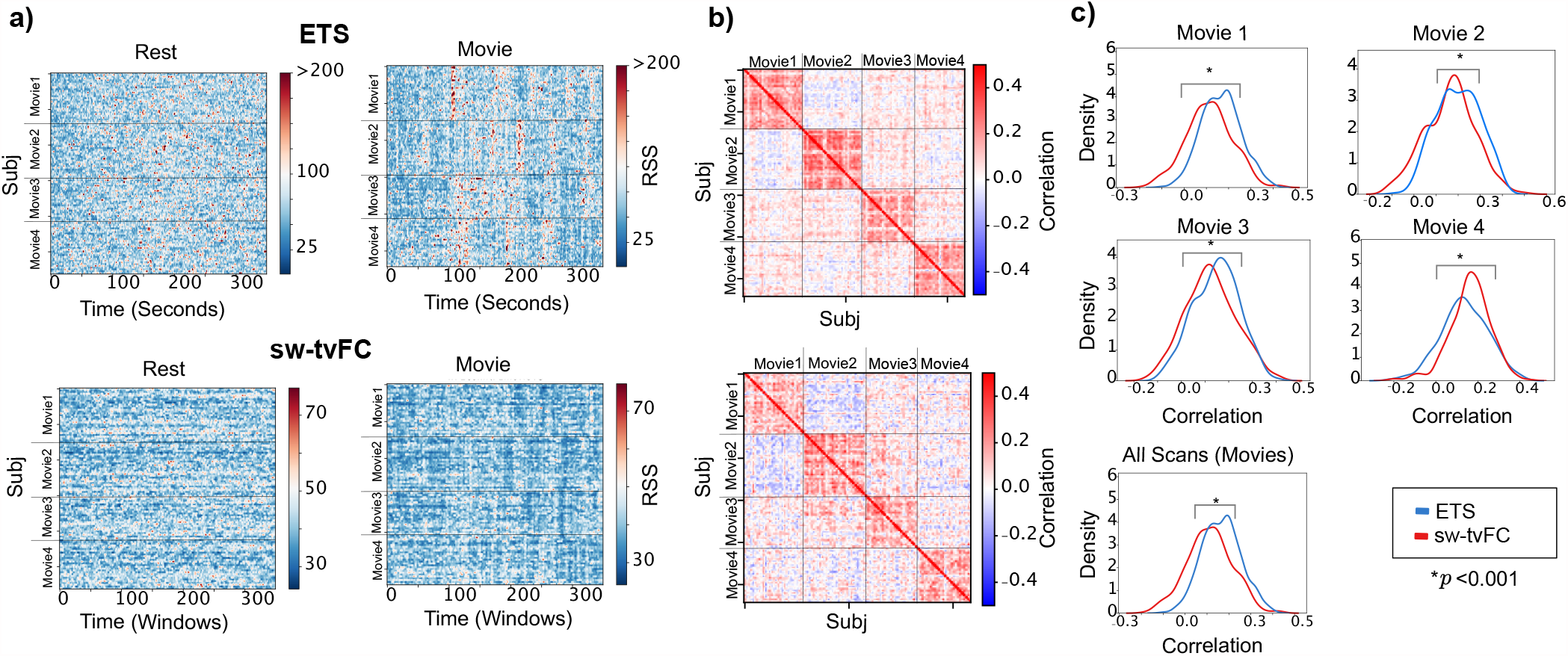
Comparison of edge time series (ETS) and sliding window-time varying functional connectivity (swtvFC) in identifying consistent co-fluctuation patterns across subjects in rest versus movie-watching conditions. *(a)* Shows the root sum square (RSS) of interpolated edge time series per subject during movie-watching and rest conditions. When comparing rest versus movie-watching conditions for both methods, RSS patterns specifically, high RSS values (shown in red colors) are consistent across subjects for ETS in movie-watching conditions. *(b)* Shows the inter-subject similarity (ISS) based on RSS pattern for movie-watching condition where ISS is higher in ETS (*p* < 0.0001). *(c)* Shows the distribution of ISS values (elements in the upper diagonal of matrices shown in panel *b*) for individual movie scans and all the movie scans together where ETS shows higher ISS compared to sw-tvFC.

#### Whole-brain co-fluctuation patterns at peaks and troughs

In the previous section, we demonstrated that ETS provide a synchronized estimation of the co-fluctuation patterns, specifically high-amplitude ones across subjects which indicates their unique contribution to the overall connectivity patterns. In this section, we further examined the role of these high-amplitude co-fluctuations and their distinction with low-amplitude ones. To do this, we defined measures of trough-to-trough duration and the peak co-fluctuation amplitude between two troughs of the RSS signal (Figure 4a) which allows evaluating the relationship between the high- and low-amplitude co-fluctuations. We found that, compared to sw-tvFC, ETS exhibited higher peaks and shorter trough-to-trough duration (*t*-test, *p* < 0.001; Figure 4c), which further indicated that ETS reflects rapid fluctuations over time. Moreover, we calculated the similarity between peaks and troughs as the correlation coefficient between whole-brain co-fluctuations at peaks and troughs. Our results suggested that there is a lower similarity between peaks and trough in terms of collective co-fluctuations in ETS than sw-tvFC (*t*-test, *p* < 0.001; Figure 4c).

**FIG. 4.**
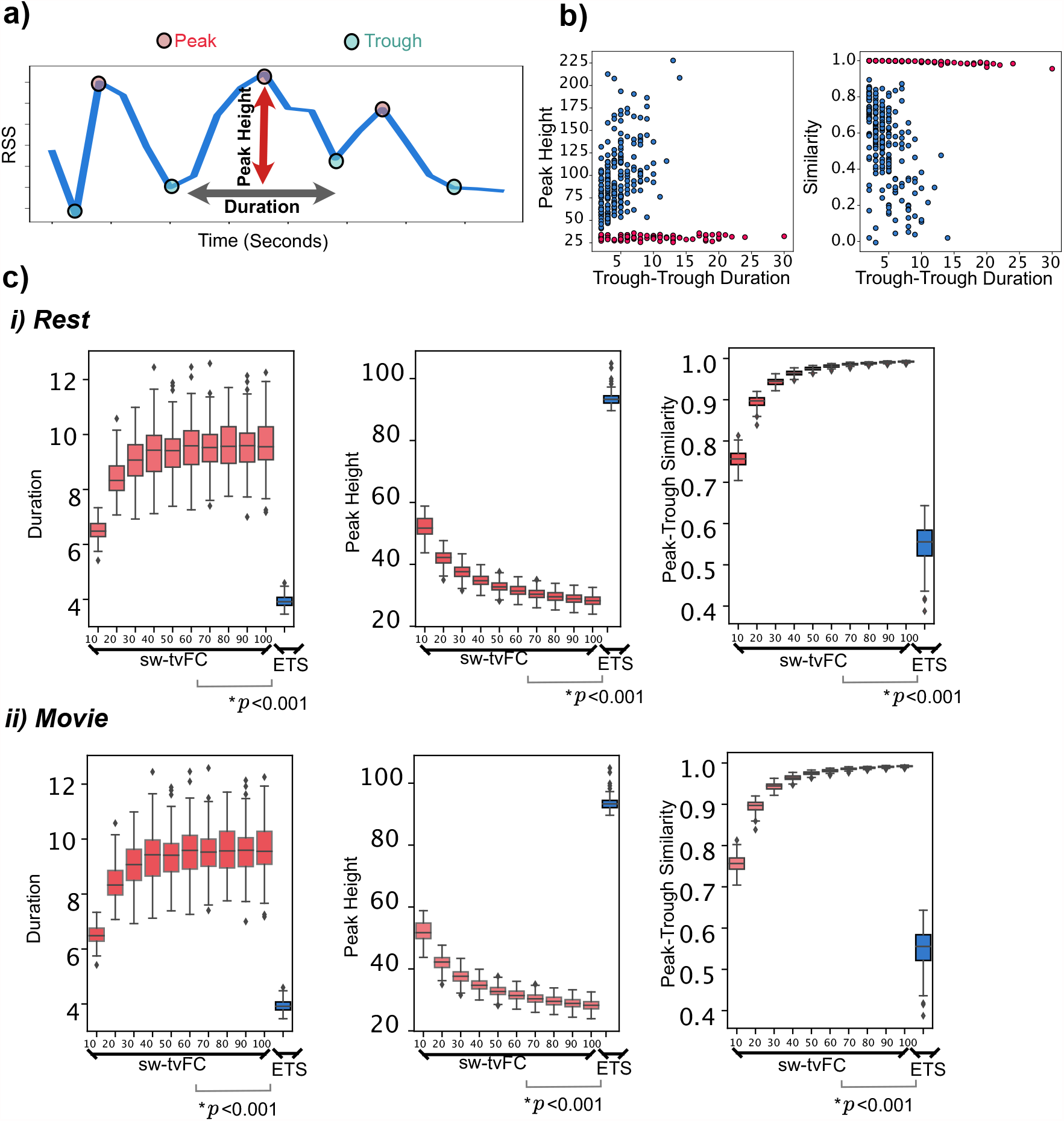
Peak-trough relationship in the whole-brain co-fluctuation patterns measured by root sum square (RSS) signal. *(a)* Shows the procedure for calculating peak amplitude and duration measures in the RSS signal. Troughs are identified as time points where their amplitude are lower than their two direct neighbors. *(b)* Shows an example of peaktrough relationship in one subject. *(c)* Comparing the peak-trough relationship in ETS and sw-tvFC (w=10-100 frames with increments of 10).

### Edge time series in autism spectrum disorder

#### Edge fluctuations in autism spectrum disorder vs. healthy controls

In the previous section, we discussed differences between ETS and sw-tvFC in terms of their ability to capture time-varying features of functional brain networks. Our findings suggested the effectiveness of ETS in tracking rapid transitions in the magnitude of collective co-fluctuations, as evidenced by greater co-fluctuation amplitude and shorter trough-to-trough duration relative to sw-tvFC. In this section, we used ETS to examine the collective, i.e. whole-brain, and edge-level co-fluctuations over time. More specifically, we used the two previously defined measures of trough-to-trough duration and the peak co-fluctuation amplitude to examine the differences of ASD and CN during passive viewing of naturalistic movies.

First, we examined differences in collective co-fluctuations of brain regions between ASD and CN in terms of trough-to-trough duration and the peak co-fluctuation amplitude. To this end, we calculated the average trough-to-trough duration and the peak amplitude of the RSS signal for each subject in both the ASD and CN groups. Our results, as shown in Figure 5a, suggested similar patterns of peak co-fluctuations between CN and ASD (*t*-test, *p* = 0.97). However, a close examination revealed subtle distinctions between the two groups. Specifically, we found that trough-to-trough duration was greater in ASD, compared to CN (*t*-test, *p* = 0.005; Figure 5a). To ensure that these differences were not driven by head motion, we conducted a posthoc motion correction analysis in which we regressed out the mean head motion (e.g., derivative of in-scanner motion and framewise displacement) from trough-to-trough duration and the peak co-fluctuation amplitude measures and compared the obtained residuals between ASD and CN. The results of this analysis were in line with the original findings, suggesting that ASD and CN are different in terms of the trough-to-trough duration (*t* test, *p* = 0.01), but not in terms of peak amplitude (*t* test, *p* = 0.35; Figure S5).

**FIG. 5.**
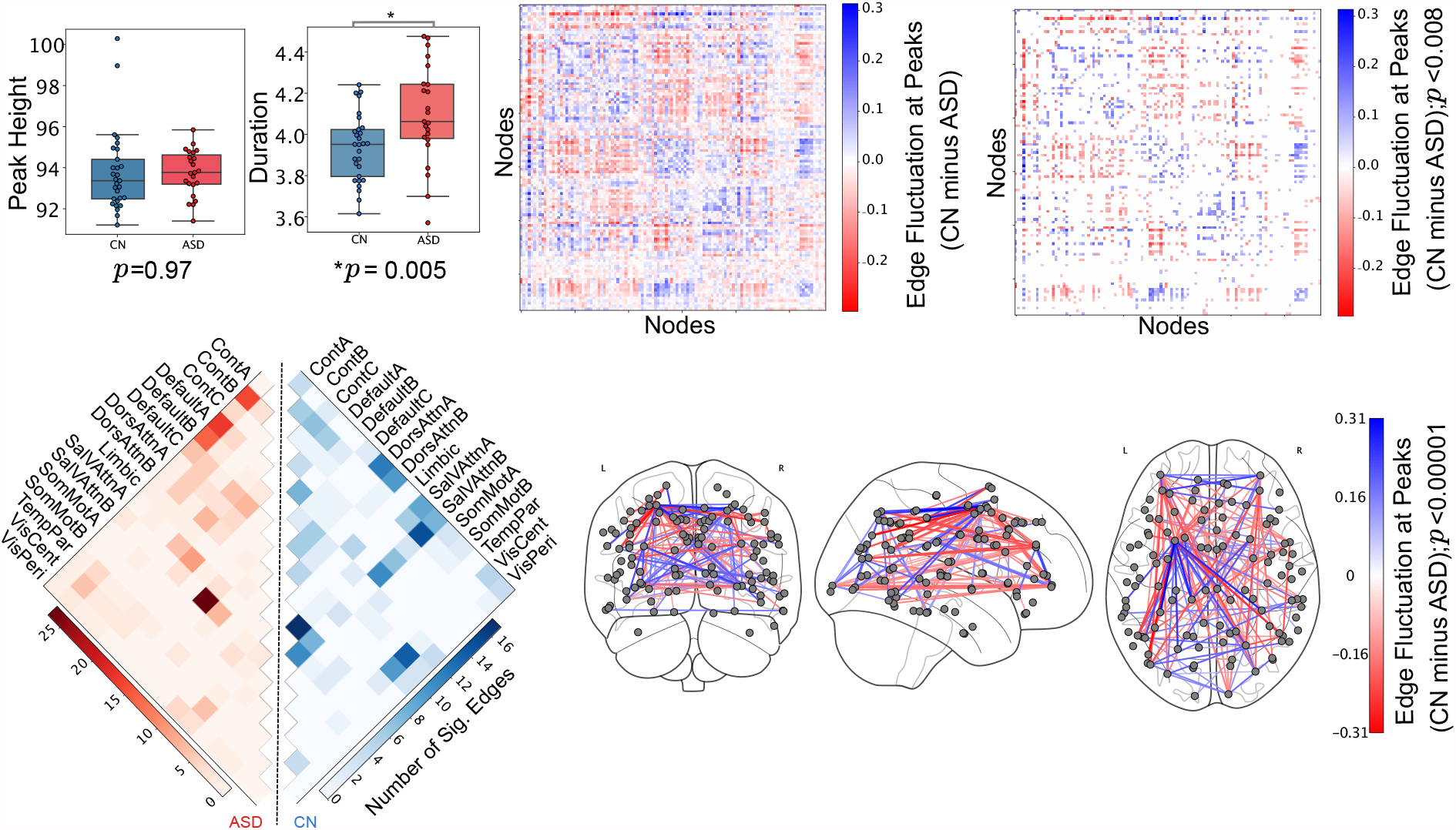
Edge time series (ETS) in autism spectrum disorder (ASD) and control (CN) subjects during movie-watching condition. *(a)* Average of peaks and trough-to-trough duration between ASD and CN group. Each point in the box plot shows the average of peaks/trough-to-trough duration measure for one subject across scans.*(b)* Averaged differences of edges in peak fluctuation amplitude between CN and ASD (CN minus ASD). *(c)* Edges that are different in peak fluctuation amplitude between ASD and CN (*p*_*adjusted*_ = 0.0075,false discovery rate (FDR)= 0.05). *(d)* Edges shown in panel c sorted based on Yeo 17 functional networks [31]. Each cell represents the number of significant edges (blue (CN > ASD), red (ASD > CN)). *(e)* Visualization of edges shows in panel c, the *p* value cutoff is selected for visualization.

Next, we conducted an edge-wise comparison of high-amplitude co-fluctuations between ASD and CN. Our results showed that the within-network edges, i.e. those that link nodes belonging to the same brain system, exhibited greater magnitude fluctuations in CN (*t*-test, *p*_*adjusted*_ < 0.008, false discovery rate (FDR) = 0.05; Figure 5b-e). The results presented in this section were generated using data from all subjects pooled across all scans. The results for individual scans are available in the supplementary section (Figures S6 and S7).

## DISCUSSION

In this paper, we compared dynamic properties of ETS with the commonly used method for estimating tvFC, sliding window. We conducted our comparisons in several steps including state transition, co-fluctuation synchrony across subjects, and so on. We found that ETS capture faster and bursty network dynamics, which is often not feasible in sw-tvFC due to the blurring effect induced by windowing procedure. Building on this important feature of ETS, we used ETS to compare co-fluctuation patterns between ASD and CN groups. We found that at the whole-brain co-fluctuation level, while CN and ASD show similar levels of peak amplitude co-fluctuations, ASD shows higher trough-to-trough duration.

### Edge time series characterize fast and bursty network dynamics

A growing number of studies have modeled time-varying changes in network structure to study fast changes in network dynamics and linking their features to inter-individual variation in traits, cognition, and clinical status [32–34]. Although many techniques are available for estimating and studying time-varying networks, the most common is the sliding window method. This approach, however, requires the user to define several key parameters, each of which impact the character of the estimated networks. Additionally, the use of a sliding window, which includes multiple successive samples, prohibits the localization of networks to a specific point in time.

However, there exists several methods that can be used to partially address this issue [33, 35, 36]. Among these methods is the recently-proposed “edge time series”. This method decomposes FC into its exact frame-wise contributions, generating an estimate of the co-fluctuation magnitude between node pairs at each point in time, thereby obviating the need for a sliding window. Although this method has been used in several papers [25–27], which have documented characteristics not usually reported in sliding window estimates of tvFC, e.g., bursts of co-activity, no direct comparison of edge time series and sliding-window tvFC has been carried out.

Our study fills this gap in the literature, measuring several commonly reported variables in both edge time series and sliding-window tvFC. We find that, broadly, these two methods yield estimates of time-varying networks that are globally similar, with the similarity peaking when sliding-window tvFC is estimated using short (but not the shortest) window duration. This location of this peak may reflect tradeoffs between ability to network reconstruction accuracy, which improves with many samples, and temporal precision, which increases with fewer samples. We also found that edge time series have a shorter “memory” than sw-tvFC, demonstrating that not only does temporal autocorrelation decay to a baseline value faster than sw-tvFC, the baseline itself is established at a lower level. Finally, we used a common clustering technique to define network “states” and calculated the probabilities of transitions from one state to another. We found that recurrent transitions were more common in sw-tvFC compared to edge time series and that transitions to other states is more common in edge times series.

These findings inform our understanding and interpretation of brain dynamics. Sliding-window estimates paint a picture in which the brain tends to slowly traverse a high-dimensional state space, with its state at *t* + 1 highly dependent on its previous state at time *t*. In contrast, edge time series exhibit faster velocities, rapidly reconfiguring over short timescales with punctuated, high-amplitude bursts. Notably, however, both techniques operate on the same input data – nodal time series. That they offer dissimilar insight highlights the potential for ETS to serve as a complementary approach to the conventional sliding-window method.

### Relevance of high-amplitude co-fluctuations to cognition and behavior

Previous studies have examined edge time series and characterized some of their basic properties [25, 26], speculating that these properties might serve as potent biomarkers for comparing individuals in terms of their cognitive or clinical states. However, with limited exceptions, these speculations have not been investigated. Here, as part of an exploratory analysis, we performed two analyses. First, we compared edge time series and sw-tvFC in terms of their ability to capture inter-subject correlations across individuals during passive viewing of movies. To this end, we found that when using ETS, whole-brain co-fluctuation patterns (RSS values across time) are more similar across subjects during movie-watching condition compared to sw-tvFC. This observation highlights a strength of ETS in capturing shared responses across subjects to the mutual stimuli. On the other hand, we also found that, by examining whole-brain connectivity profiles during peaks and troughs, the similarity between peaks and troughs was lower using ETS compared to sw-tvFC. Collectively, these results suggest that the temporal precision afforded by edge time series may allow us to better track *when* brains respond to stimuli, while exposing heterogeneity of response profiles (connectivity patterns) across individuals. We speculate that these two features could be taken advantage of by future studies investigating brain-behavior relationships.

### Edge time series disclose differences between ASD and healthy control dynamics

Another key finding of this paper is that ASD, compared to CN, shows longer trough-to-trough duration, but similar peak amplitudes in the whole-brain co-fluctuation patterns (RSS signal) during movie-watching. This observation suggests that, although ASD patients respond similarly to stimuli as controls, their network dynamics are systematically “stickier” than those of controls – taking longer to rise to peak activity and then return to baseline. These stickier dynamics may have important implications for the understanding of disorders and disease. For instance, slower dynamics could impede or delay transitions between brain states and, to the extent that brain states are of cognitive relevance, could impact the timing of ongoing cognitive processes [37–40].

More generally, these observations underscore the possibility that population-level differences, if they exist, may be encoded not in the structure of a network, but in its dynamics and changes across time. Indeed, a growing number of studies have shown that features such as transition rate and occupancy time of dynamic network states vary with age and differ between clinical conditions [41–43]. Higher-order network structure, including its system- and module-level architecture, also vary across time, and has been shown in previous studies to track with individual differences in a variety of measures [44, 45].

## LIMITATIONS AND FUTURE WORK

In this work, we compared sliding window and ETS methods for estimating tvFC, further using ETS to investigate inter-subject correlations during movie-watching and group differences between ASD and control populations. Although the results of this paper help contextualize ETS with respect to existing methods for estimating tvFC and highlight its potential as a method for studying inter-individual differences, it has a number of limitations. At the same time, it presents exciting opportunities for future work.

One way to broaden our findings is to extend the analysis of network states reported in the first part of the paper and compare the control and ASD groups in terms of these states. Previous studies have shown that these states undergo individualization and may present useful and subject-specific information for comparing groups [26]. Additionally, the framework applied here to an ASD population could be extended to other clinical populations. Indeed, there exist many large, publicly available datasets that include both clinical groups [46] and massive control populations that are accompanied by sub-clinical responses to assessments of different neuropsychiatric disorders [47, 48].

Another possible extension includes exploring edge functional connectivity (eFC), which refers to the correlation structure of edge time series. Previous studies demonstrated that this higher-order construct is both highly reliable and can readily identify overlapping communities in networks. Yet another opportunity for future work includes more detailed benchmarking of ETS using synthetic example, with the aim of clearly distinguishing features that are genuinely “dynamic” from those can be explained by time-invariant features of the static FC matrix [49]. While our work clearly demonstrates that it returns dissimilar results relative to sliding window methods, it remains unclear whether those dissimilarities necessarily mean that ETS is outperforming the other approach.

Finally, while our work demonstrates that there are systematic differences in trough-to-trough duration and, possibly, the height of peaks, it does not speak to when those differences occur. Nor does it speak to the character of the stimulus present at those instants. Future work using annotated naturalistic stimuli could be undertaken to help address these questions.

## MATERIALS AND METHODS

### Dataset

We analyzed fMRI data of 29 CN and 23 ASD individuals that are scanned multiple times during resting-state and movie-watching conditions. The number of subjects used in this study for scan 1, 2, 3 and 4 were subsequently 29 CN, 23 ASD; 29 CN, 23 ASD; 26 CN, 20 ASD; and 25 CN, 21 ASD. The details for this dataset including participant characteristics, data acquisition, and preprocessing pipeline can be found in [29].

### Image Preprocessing

#### MRI acquisition and processing

MRI images were acquired using a 3T whole-body MRI system (Magnetom Tim Trio, Siemens Medical Solutions, Natick, MA) with a 32-channel head receive array. Both raw and prescan-normalized images were acquired; raw images were used at all preprocessing stages and in all analyses unless specifically noted. During functional scans, T2*-weighted multiband echo planar imaging (EPI) data were acquired using the following parameters: TR/TE = 813/28 ms; 1200 vol; flip angle = 60 ; 3.4 mm isotropic voxels; 42 slices acquired with interleaved order covering the whole brain; multi-band acceleration factor of 3. Preceding the first functional scan, gradient-echo EPI images were acquired in opposite phase-encoding directions (10 images each with P-A and A-P phase encoding) with identical geometry to the EPI data (TR/TE = 1175/39.2 ms, flip angle = 60) to be used to generate a fieldmap to correct EPI distortions, similar to the approach used by the Human Connectome Project [50]. High-resolution T1-weighted images of the whole brain (MPRAGE, 0.7 mm isotropic voxel size; TR/TE/TI = 2499/2.3/1000 ms) were acquired as anatomical references. All functional data were processed according to an in-house pipeline using FEAT (v6.00) and MELODIC (v3.14) within FSL (v. 5.0.9; FMRIB’s Software Library, www.fmrib.ox.ac.uk/fsl), Advanced Normalization Tools (ANTs; v2.1.0) [51], and Matlab R2014b. This pipeline was identical to the GLM + MGTR procedure described in [52].

In more detail, individual anatomical images were bias corrected and skull-stripped using ANTs, and segmented into gray matter, white matter, and CSF partial volume estimates using FSL FAST. A midspace template was constructed using ANTs’ buildtemplateparallel and subsequently skull-stripped. Composite (affine and diffeomorphic) transforms warping each individual anatomical image to this midspace template, and warping the midspace template to the Montreal Neurological Institute MNI152 1mm reference template, were obtained using ANTs.

For each functional run, the first five volumes (≃4 seconds) were discarded to minimize magnetization equilibration effects. Framewise displacement traces for this raw (trimmed) data were computed using fsl motion outliers. Following [29, 53], we performed FIX followed by mean cortical signal regression. This procedure included rigid-body motion correction, fieldmapbased geometric distortion correction, and non-brain removal (but not slice-timing correction due to fast TR [50]). Preprocessing included weak highpass temporal filtering (> 2000 s FWHM) to remove slow drifts [50] and no spatial smoothing. Off-resonance geometric distortions in EPI data were corrected using a fieldmap derived from two gradient-echo EPI images collected in opposite phase-encoding directions (posterior-anterior and anterior-posterior) using FSL topup.

We then used FSL-FIX [54] to regress out independent components classified as noise using a classifier trained on independent but similar data and validated on handclassified functional runs. The residuals were regarded as “cleaned” data. Finally, we regressed out the mean cortical signal (mean BOLD signal across gray matter partial volume estimate obtained from FSL FAST). All analyses were carried out on these data, which were registered to subjects’ skull-stripped T1-weighted anatomical imaging using Boundary-Based Registration (BBR) with epi reg within FSL. Subjects’ functional images were then transformed to the MNI152 reference in a single step, using ANTS to apply a concatenation of the affine transformation matrix with the composite (affine + diffeomorphic) transforms between a subject’s anatomical image, the midspace template, and the MNI152 reference. Prior to network analysis, we extracted mean regional time series from regions of interest defined as sub-divisions of the 17-system parcellation reported in [31] and used previously [55–57]. Wakefulness during movie and rest scans was monitored in real-time using an eye tracking camera (Eyelink 1000).

#### Naturalistic stimuli

All movies were obtained from Vimeo (https://vimeo.com). They were selected based on multiple criteria. First, to ensure that movies represented novel stimuli, we excluded any movie that had a wide theatrical release. Secondly, we excluded movies with potentially objectionable content including nudity, swearing, drug use, etc. Lastly, we excluded movies with intentionally startling events that could lead to excessive in-scanner movement.

Each movie lasted approximately 1 to 5 minutes. Each movie scan comprised between four and six movies with genres that included documentaries, dramas, comedies, sports, mystery, and adventure. See Table S1 for more details.

### Edge (Co-fluctuation) time series

Functional brain networks are constructed by estimating the statistical dependency between fMRI BOLD activity of brain regions. The magnitude of these dependencies reflects the strength of functional connection between brain regions. One of the most common measures to estimate the dependency between brain regions is the Pearson correlation coefficient. The overall procedure for calculating Pearson coefficient is as follows: Let *x*_*i*_ = [*x*_*i*_(1), …, *x*_*i*_(*T*)] and *x*_*j*_ = [*x*_*j*_(1), …, *x*_*j*_(*T*)] be the time series recorded from voxels or parcels *i* and *j*, respectively. We can calculate the correlation of *i* and *j* by first z-scoring each time series, such that 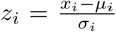, where 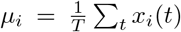 and 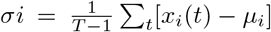 are the time averaged mean and standard deviation. Then, the correlation of *i* with *j* can be calculated as 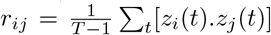. Repeating this procedure for all pairs of parcels results in a node-by-node correlation matrix, i.e., an estimate of FC. If there are N nodes, this matrix has dimensions [*N* × *N*]. To estimate edge-centric networks, we modify the above approach such that we only calculate the element-wise product of two time series and remove the step for calculating the mean. This operation would result in a vector of length *T* whose elements encode the moment-by-moment co-fluctuation magnitude of parcels *i* and *j*. More specifically, the positive values in the vector reflect the simultaneous increase or decrease in the activity of parcels *i* and *j*, while negative values reflect the opposite direction (one increasing while the other decreasing and vice versa) of the magnitude of their activity. Similarly, if either *i* or *j* increased or decreased while the activity of the other was close to baseline, the corresponding entry would be close to zero. An analogous vector can easily be calculated for every pair of parcels (network nodes), resulting in a set of edge time series. With *N* parcels, this results in 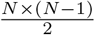 pairs, each of length *T*.

### Sliding window time series

To estimate tvFC using sliding window method, we divided every fMRI BOLD time series into several consecutive equal-sized segments (windows) and calculated correlation coefficient between time points within each window. We repeated this procedure for every window and for all pairs of time series. This results in a 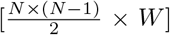 where *N* is the total number of brain regions and *W* is the total number of windows used to for sw-tvFC estimation (for every time series). We have normalized obtained sw-tvFC values (i.e., *r*) using Fisher transform 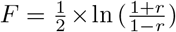. We used window sizes (*w*) of 10-100 with increments of 10; and offset (value for shifting window) = 1 to estimate sw-tvFC where the results for *w* = 20 are in provided in the main text and results for other window sizes are available in the supplementary section.

The obtained tvFC based on the sliding window is shorter than the actual fMRI BOLD time series, while ETS is exactly the same length as fMRI BOLD time series. Therefore, in order to compare the whole-brain co-fluctuation and inter-subject similarity between sw-tvFC and ETS, we used linear interpolation technique to resample (i.e., 350 time points) time series and calculate the similarity between the two interpolated time series.

### Autocorrelation

For every subject, we calculated the autocorrelation (i.e., lag=100) in ETS/sw-tvFC as the similarity of whole-brain co-fluctuation patterns at time *t* with the patterns at times *t* + 1, *t* + 2, …, *t* + 99, *t* + 100. We compared the averaged autocorrelation across subjects in ETS and sw-tvFC.

### K-means clustering and state transitions

We have used k-means clustering algorithm with Euclidean distance to cluster ETS/sw-tvFC. More specifically, we clustered time points based on the similarity of whole-brain co-fluctuation patterns at a given time point. For every subject, we obtained a clustered time series (1 × *T*) where every element represents a cluster index (i.e., brain state) at that given time point. After obtaining the clustered edge time series, we quantified the number of transitions between/within states over time. We used *k* =5, 10, and 15 as the initial number of clusters where results for *k* = 5 is provided in the main text and *k* = 10 and *k* = 15 are provided in the supplementary section.

### Trough-to-trough duration and peak amplitude measures

For every subject, we calculated the root sum square (RSS) of all the edge time series at every given time point resulting in a single time series. Next, we identified troughs in RSS signal and defined two measures of peak amplitude (highest peak between two troughs) and duration of trough-to-trough. Troughs (here, referring local minima) in RSS signal were defined as time points where their values were lower than the amplitude of their two direct neighbors. We used the mean peak amplitudes and trough-to-trough duration in RSS signal to compare ETS and sw-tvFC across subjects. The same approach was used to compare CN and ASD in terms of these measures.

### Correlation between confounding variables and tvFC

We conducted a posthoc motion correction analysis to examine the effect of head motion and noise in calculating trough-trough duration and peak amplitude measures in RSS signal. For every subject, we regressed out the mean of two head motion variables (e.g., derivative of scanner motion and framewise displacement) from trough-to-trough duration and the peak co-fluctuation amplitude measures and compared the obtained residuals between ASD and CN. More specifically, we regressed out the mean of head motion variables from peak amplitude measures at time points corresponding to peaks. For trough-to-trough duration measure, we took the average of head motions between every two troughs and regressed out those from the trough-trough duration measure. Finally, we compared the obtained residuals between ASD and CN groups.

## ACKNOWLEDGMENTS

This material is based upon work supported by the National Science Foundation under Grant No. 076059-00003C (RFB, OS, FZE). This research was supported by the Indiana University Office of the Vice President for Research Emerging Area of Research Initiative, Learning: Brains, Machines and Children (F.Z.E. and R.F.B.). This work was supported by the NIH (grants R01MH110630 and R00MH094409 to D.P.K. and T32HD007475 Postdoctoral Traineeship to L.B.).

---

**FIG. S1.**
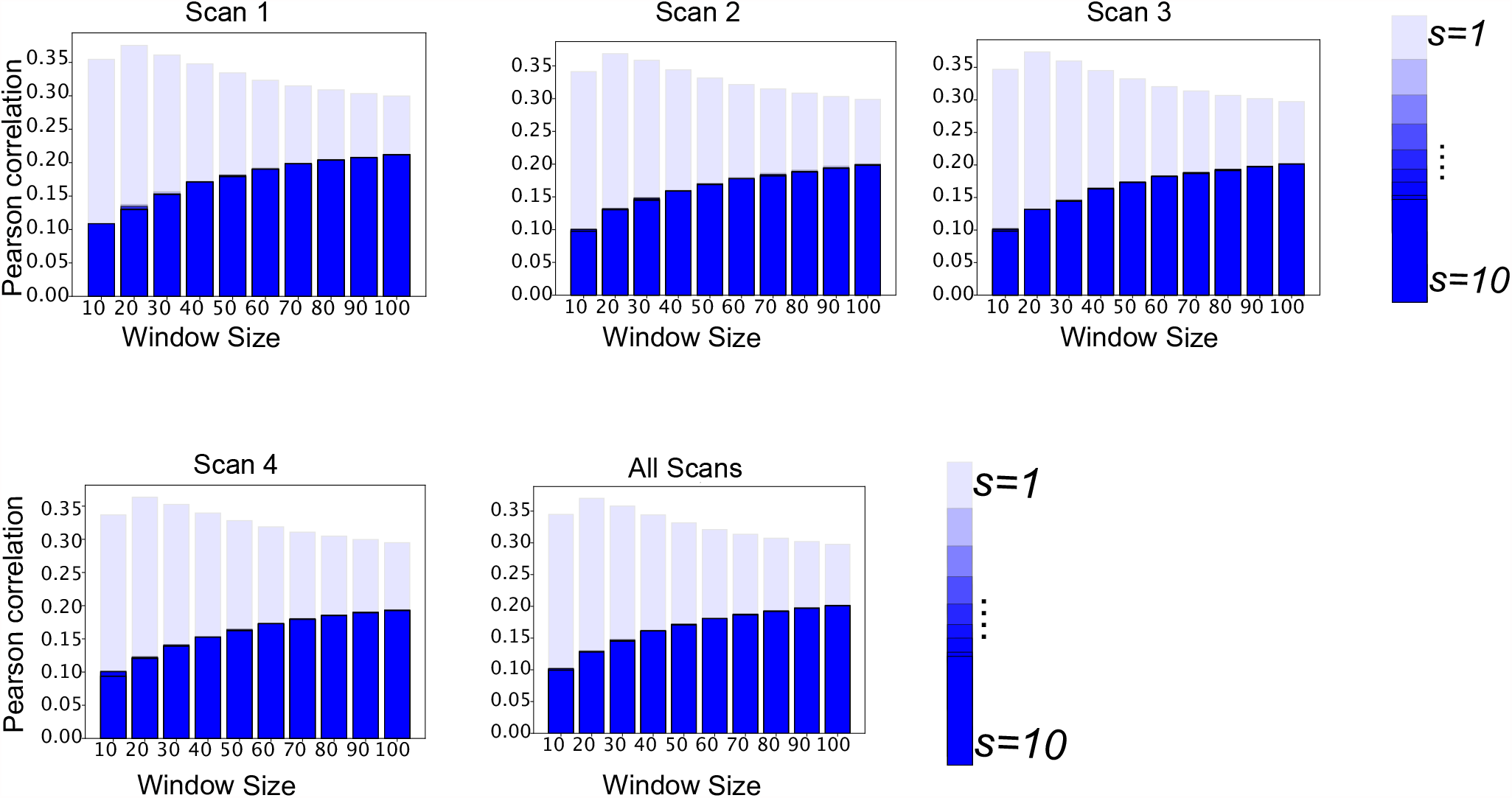
Relationship between edge time series (ETS) and sliding window time varying functional connectivity (sw-tvFC) for individual/all scans in the rest condition. Bar plots represent the averaged Pearson correlation between all the edges in edge time series and sw-tvFC across subjects in rest condition (separate scans and all scans pooled together). Shade of the colors represent the size for shifting moving window (s). For s=1, ETS and sw-tvFC are most similar for window size = 20, however, for other values of s, the similarity increases as the window size increases.

**TABLE S1.** Movie trailers included in each movie scan.

| Scan | Title | Genre | Runtime |
| --- | --- | --- | --- |
| 1 | Man Up and Go | documentary/emotional | 4m20s |
| 1 | The First 70 | documentary | 3m |
| 1 | Fixation | documentary/adventure | 1m42s |
| 1 | The Living | drama | 2m |
| 1 | SAMSARA | documentary/“unparalleled sensory experience” | 1m35s |
| 1 | Blood Brother | documentary | 2m20s |
| 2 | Birdmen | documentary/adventure | 3m59s |
| 2 | Groomed | drama | 1m30s |
| 2 | Cold | outdoor/sports | 2m |
| 2 | Sleepwalkers | drama | 2m |
| 2 | A Kind of Show | comedy | 1m |
| 3 | Geofish | documentary/adventure | 4m40s |
| 3 | The Debut | outdoor/sports | 3m23s |
| 3 | Dreams of a Life | documentary/mystery | 2m10s |
| 3 | The Front Man | documentary | 2m30s |
| 3 | This Is Vanity | drama | 1m |
| 4 | Planetary | documentary | 4m30s |
| 4 | Sign Painters | documentary | 2m50s |
| 4 | Florida Man | documentary/drama | 2m |
| 4 | The Sleeping Bear | drama | 3m40 |

**FIG. S2.**
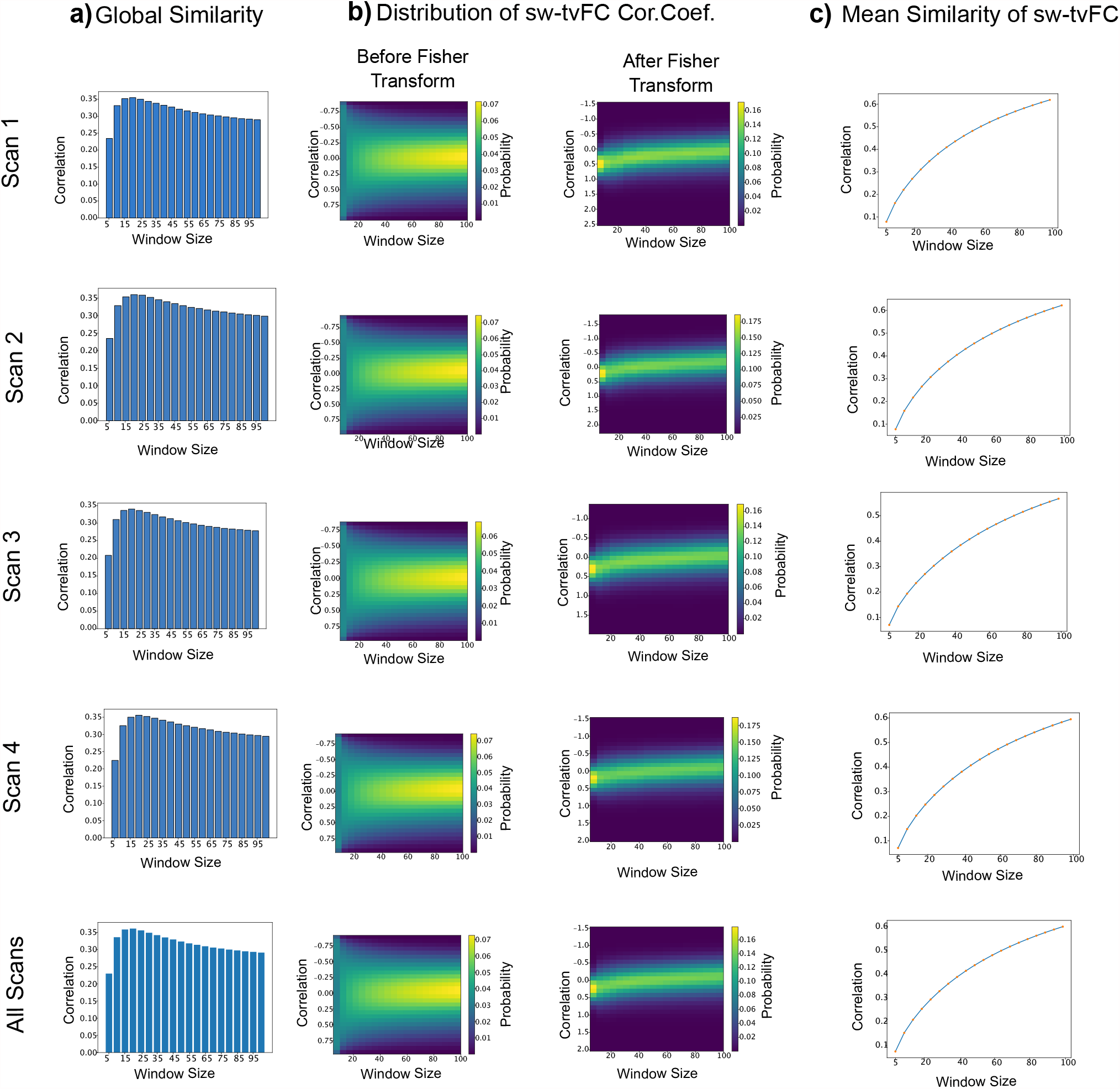
Global relationship between edge time series (ETS) and sliding window time varying functional connectivity (sw-tvFC) at rest condition. To better understand why the global correlation between ETS and sw-tvFC was not stronger, we conducted a more detailed examination, focusing on the role of window length. We hypothesized that two distinct and competing factors caused the peak correlation to occur at an intermediate window size. Specifically, we hypothesized that when the window size was very small, the sw-tvFC would be able to capture fast fluctuations in connectivity, but due to a relatively small number of samples the estimate of connection weights would be inaccurate. Conversely, longer windows provide more accurate estimates of connection weights but at the expense of temporal specificity. To test this, we systematically varied the duration of windows and found that, for very short windows, the histogram of connection weights was highly bimodal for all frames. This is in contrast to typical connection weights of ETS, which are unimodal and generally centered around zero. This mismatch of distributions likely explains why, for short windows, ETS and sw-tvFC exhibit a poor correspondence. On the other hand, as increased the length of windows, the estimated networks exhibited little variation across time, suggesting they are incapable of capturing the “burst” dynamics observed in ETS. Collectively, these results explain both the overall weak correspondence between sw-tvFC and ETS at the global scale and why the peak similarity occurs at an intermediate window size. (*a*) Global similarity of ETS and sw-tvFC (Pearson correlation coefficient) based on whole-brain co-fluctuations. (*b*) Histogram of sw-tvFC connection weights (without Fisher transforming) where for very small windows, the histogram of connection weights is bimodal and dissimilar from that of ETS. After normalizing sw-tvFC weights using Fisher transforming, the histogram of connection weights leads to a distribution with a single peak, but with high levels of variability. (*c*) Average similarity of sw-tvFC at time t to all time points where for very large windows, the average similarity becomes very large, meaning that temporal specificity is also likely reduced.

**FIG. S3.**
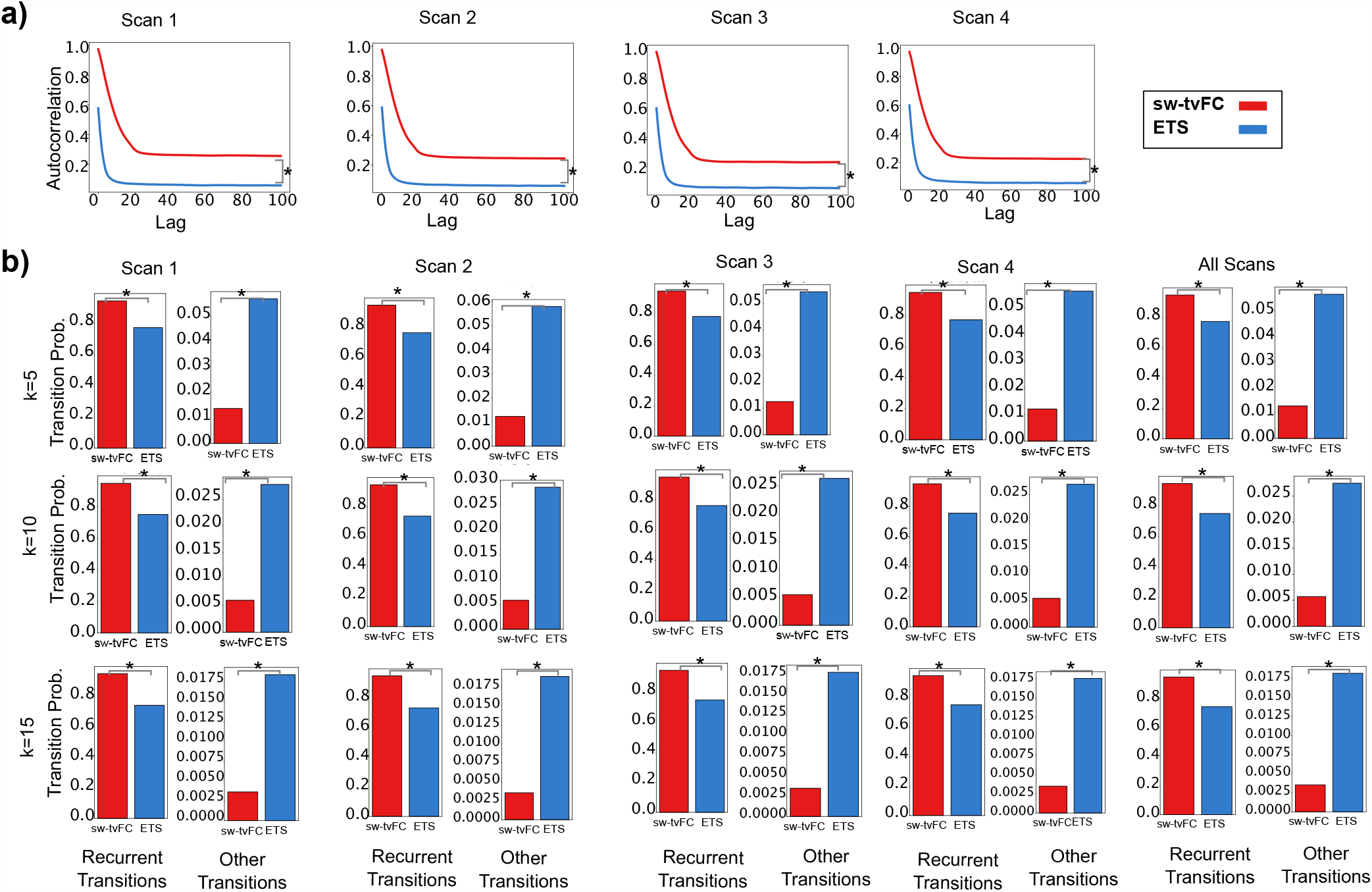
Autocorrelation and state transition probabilities of edge time series (ETS) and sliding window time varying functional connectivity (sw-tvFC) for individual scans in the rest condition. (*a*) Shows the autocorrelation in ETS and sw-tvFC (i.e., window size (*w* = 20)). ETS shows lower rate of autocorrelation, suggesting the presence of high-amplitude co-fluctuations (*t* test, *p* < 0.001). This was also evident in the state transition plot (*b*), where ETS have higher between-state transition and lower within-state transition compared to the sw-tvFC (*t* test, *p* < 0.001). States at each given time point were defined based on the whole-brain co-fluctuations using *k*-means clustering algorithm (*k*= number of initial clusters).

**FIG. S4.**
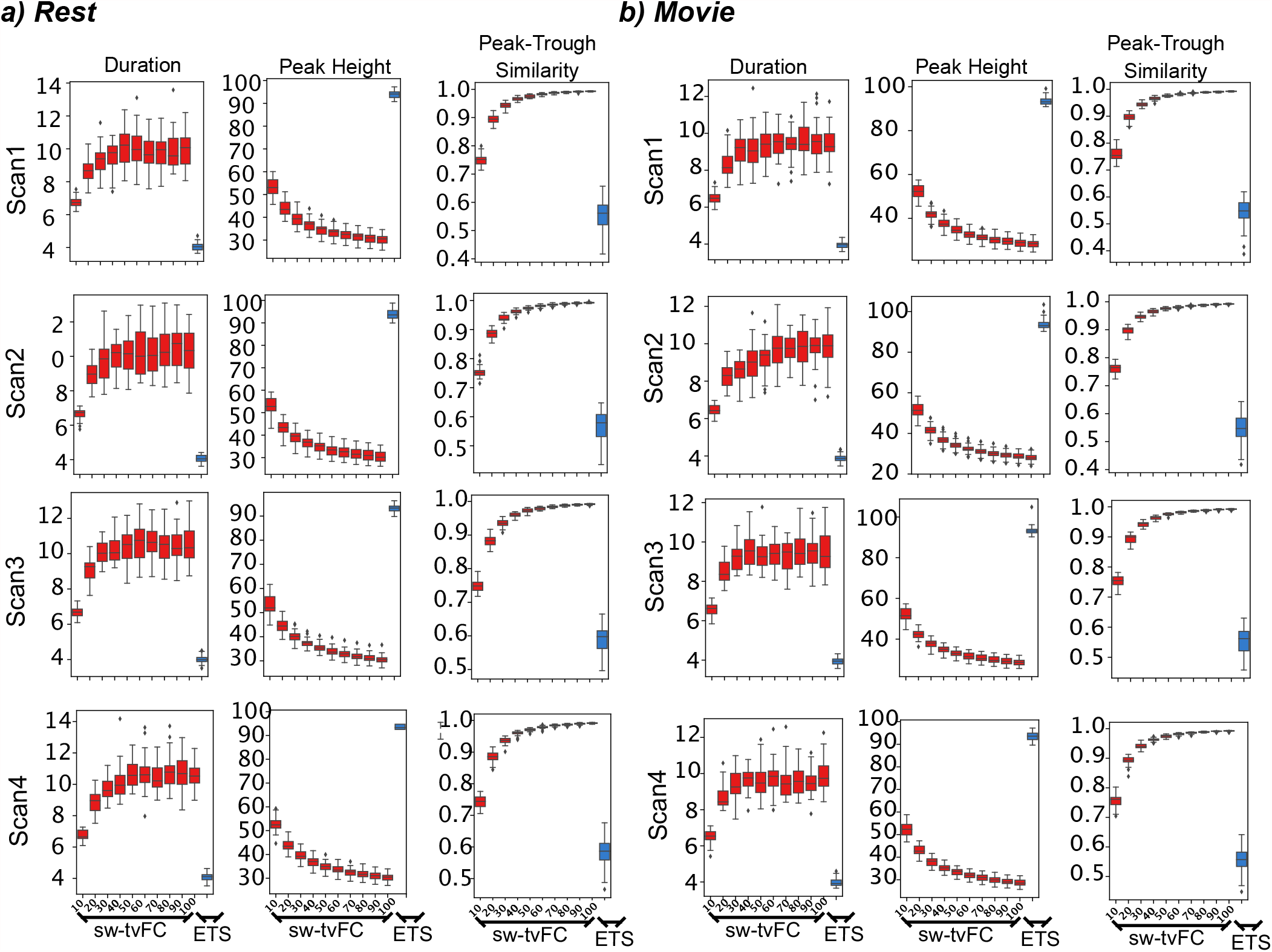
Peak-trough relationship in the whole-brain co-fluctuation patterns in rest and passive movie-watching conditions (individual scans). *(a-b)* Compares the peak-trough relationship in edge time series versus sliding window time varying functional connectivity (w=10-100 frames, with an increment of 10) in rest and passive movie-watching conditions subsequently (*t* test, *p* < 0.001).

**FIG. S5.**
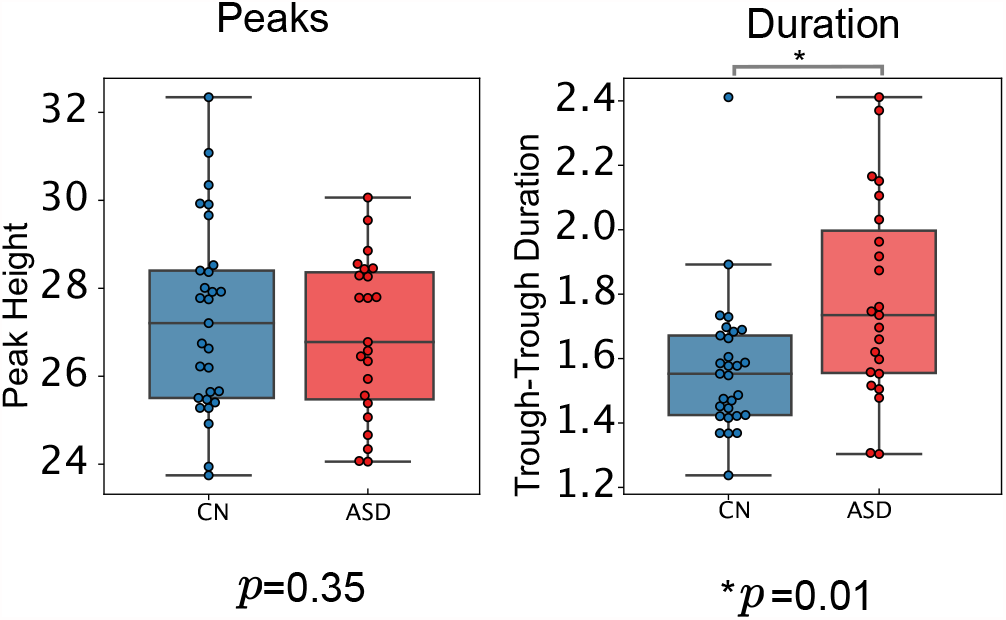
Effect of head motion on measuring the collective co-fluctuations of brain regions during passive movie-watching condition using edge time series (ETS). We conducted a posthoc motion correction analysis in which we regressed out the mean of head motion variables (e.g., derivative of scanner motion and framewise displacement) from trough-to-trough duration and the peak co-fluctuation amplitude measures and compared the obtained residuals between autism spectrum disorder (ASD) and control (CN) subjects. Each point in the box plot shows the average residuals of peaks/trough-to-trough duration measure for one subject across scans. These results are in line with the original findings, suggesting that ASD and CN are different in terms of the trough-to-trough duration (*t*-test, *p* = 0.01), but not in terms of peak amplitude (*t*-test, *p* = 0.2)

**FIG. S6.**
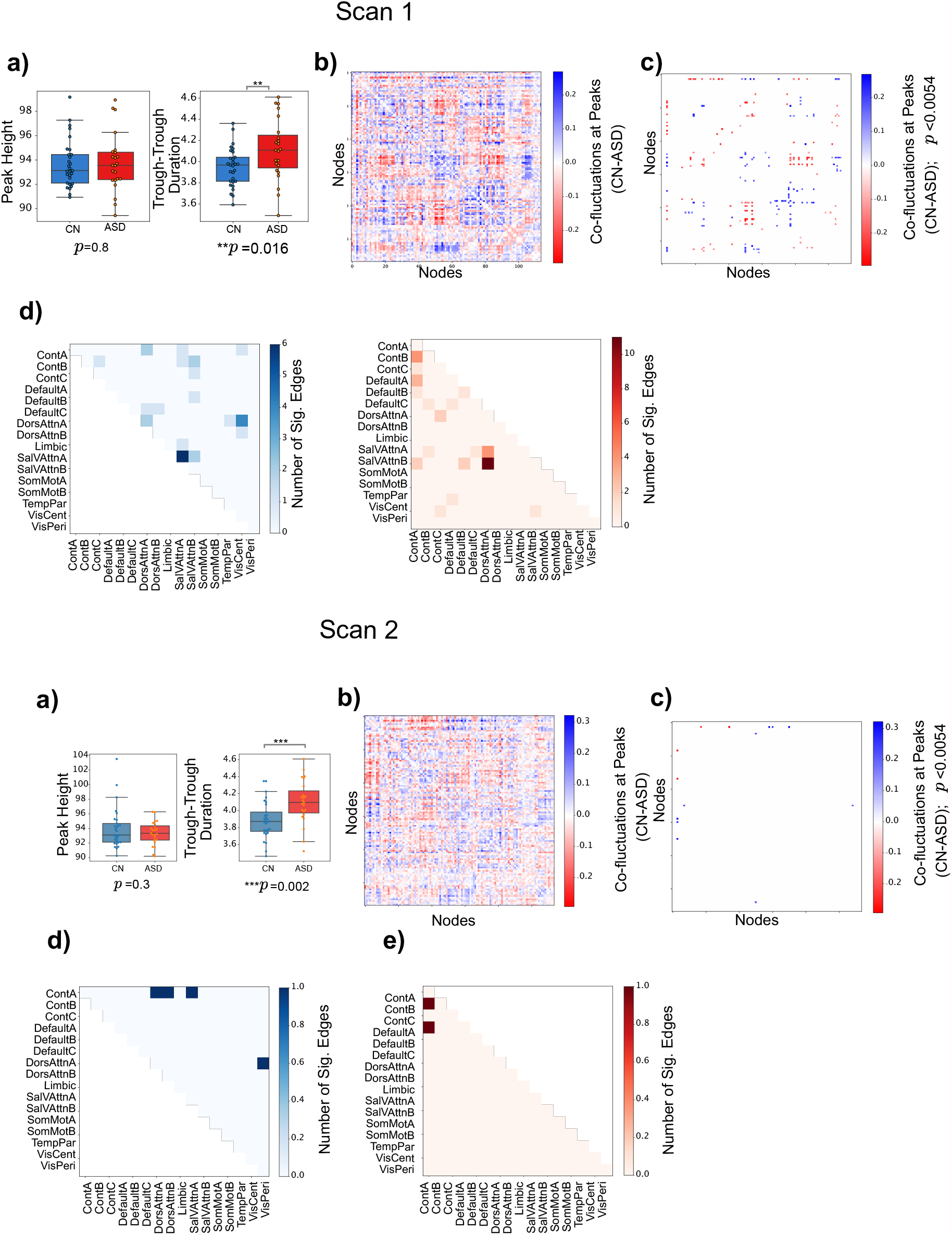
Edge time series in autism spectrum disorder (ASD) and healthy controls (CN) during passive movie-watching condition (individual scans). *(a)* Peaks and trough-to-trough duration calculated for the ASD and CN subjects. (*b*) Averaged differences of edges in peak fluctuation amplitude between CN and ASD (CN minus ASD). *(c)* Edges that are different in peak fluctuation amplitude between ASD and CN (for scans 1 and 2; *p*_*adjusted*_ = (0.0054, 0.00034),false discovery rate= 0.3). *(d,e)* Edges shown in panel *c* sorted based on Yeo 17 functional networks [31]. Each cell represents the number of significant edges (blue (CN>ASD), red (ASD>CN)).

**FIG. S7.**
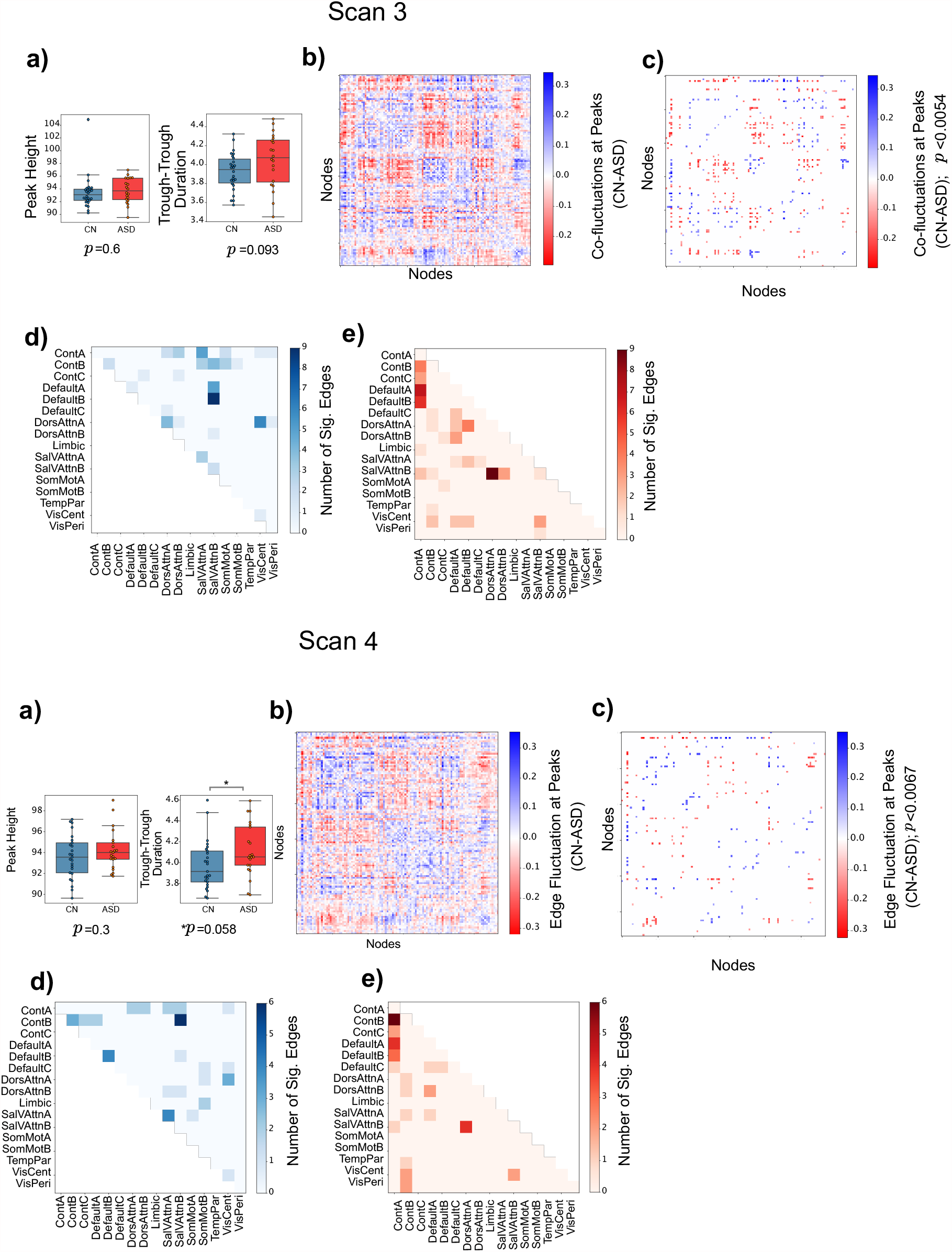
Edge time series in autism spectrum disorder (ASD) and healthy controls (CN) during movie-watching condition (individual scans). *(a)* Peaks and trough-to-trough duration calculated for the ASD and CN subjects. *(b)* Averaged differences of edges in peak fluctuation amplitude between CN and ASD (CN minus ASD). *(c)* Edges that are different in peak fluctuation amplitude between ASD and CN (for scans 3 and 4; *p*_*adjusted*_ = (0.012, 0.0067),false discovery rate= 0.3). *(d,e)* Edges shown in panel *c* sorted based on Yeo 17 functional networks [31]. Each cell represents the number of significant edges (blue (CN>ASD), red (ASD>CN)).

